# The Application of Label-free Detection Using a Tapered Optical Fiber System for Head and Neck Cancer and Infectious Biomolecules

**DOI:** 10.1101/2023.09.25.559133

**Authors:** Casey Collet, Cong Deng, Chaminda Ranathunga, Partha P. Banerjee, De-Chen Lin, Uttam Sinha

## Abstract

**Introduction:** Head and neck squamous cell carcinoma (HNSCC) is associated with high morbidity and mortality due to late detection. Tapered optical fiber sensors (TOFS) are biosensors with the potential application as a point-of-care device for detection of HNSCC biomarkers. TOFS uses optical fibers as transduction elements and antigen-antibody binding for the detection of target biomolecules. The present TOFS system was designed to achieve high specificity, sensitivity, and repeatability. To explore its application in HNSCC, we targeted the proinflammatory cytokine IL-8, known for its role in promoting tumorigenicity, metastasis, and angiogenesis in HNSCC. To validate our proof-of-concept experiment, a viral surrogate of SARS-CoV-2, Human Coronavirus OC43 (HCoV-OC43), was also tested.

**Methods:** Our TOFS system contains four main parts: the tapered optical fiber, reservoir (cell), laser light source, and photodetector. The fiber is bitapered with a narrow waist region that anchors antibodies to detect target biomolecules. Light is transduced by the laser to the fiber where it travels through the down-taper region to the tapered waist followed by the up-tapered region and ultimately transmitted to the photodetector. The tapered waist facilitates the transformation of a single mode light to multiple modes, which reverts to a single mode of light as it continues through the up-taper region. Biomolecules introduced to the waist region alter the effective refractive index, which is reflected in the phase change of the wavelength. This is detected by the photodetector and quantified using Fouier analysis. Our tapered fibers were designed with antigen-antibody complexes targeting IL-8 and HCoV-OC43 to validate our TOFS design across two different antigens.

**Results:** TOFS with tethered mouse anti-human IgG (28nm) bound to dissolved IL-8 (14 nm) produces a measurable phase shift. As little as 10pg/ml or 7.1X10^5^ IL-8 molecules/μl was detected by the prototype device and the phase change represented in real-time the binding dynamic of IL-8 to the tethered IgG. To validate our proof-of-concept experiment, HCoV-OC43 dissolved in saliva was detected with a sensitivity of 50 viruses/mL.

**Conclusions:** TOFS device is a highly sensitive system capable of detecting proteins, viruses, and other biomolecules. Selectivity of the system is guaranteed by specific antigen-antibody binding, supported by the detection of IL-8 and HCoV-OC43 at the femtomolar level. The long-term goal for TOFS is its application as a point-of-care device used for detection, monitoring, and surveillance of HNSCC as well as detection of other pathogens in the clinical setting.

## Introduction

Head and neck squamous cell carcinoma (HNSCC) is the sixth most common cancer in the United States and one of the most common malignancies worldwide with 800,000 newly diagnosed cases annually.^1,2^ HNSCC includes cancers of the oral cavity, pharynx, and larynx. Risk factors include tobacco, alcohol, HPV, poor oral hygiene, old age, male sex, dietary deficiencies, ultraviolet radiation, and betel-quid chewing.^3,4^ Surgery and radiotherapy are the gold standards of treatment, often associated with postoperative functional and physical defects due to the location and advanced stage of the disease at the time of detection.

Affected patients suffer from high morbidity and mortality, with an average 5-year survival rate of less than 50%.^3^ Mortality rates remain stably high despite recent treatment advances, largely attributable to late disease detection.^3^ HNSCC are often preceded by precursor lesions such as leukoplakia, erythroplakia, oral submucous fibrosis and oral lichen planus, which often go undiagnosed due to their asymptomatic presentation.^3,4^ Early detection and intervention is critical for improved prognosis. Advancements in the field of biosensing may offer crucial technology for detection of biomarkers of HNSCC. Tapered optical fiber sensors (TOFS) are one such biosensor with potential application as a point-of-care device that could pave the way for novel HNSCC screening, treatment monitoring, and disease recurrence in the clinical setting.

TOFS employs optical fibers as transduction elements for detecting target biomolecules, including proteins, enzymes, and oligonucleotides.^5^ Benefits of TOFS as a biosensor include stability and its chemical-inertness, modifiable biosensing surface, low cost, ready availability, and potential for remote sensing.^5^ Though invented several decades ago, practical applications of TOFS devices have been difficult due to the typical tradeoff between sensitivity and reliability.^5-10^ This is due to challenges with complex data analysis for detecting an unstable electromagnetic (EM) field. The present TOFS system tackles this concern through the unique properties of our tapered optical fiber submerged in the liquid flow cell and our innovative Fourier analysis that addresses previous data analysis failures.

To explore the application of TOFS in HNSCC, the present paper targets the proinflammatory cytokine interleukin-8 (IL-8). IL-8 promotes tumorigenicity, metastasis, angiogenesis, and epithelial–mesenchymal transition (EMT) in HNSCC.^11^ Research supports the detection of IL-8 as a potential biomarker for HNSCC in both blood and saliva.^11^ To support the generalizability of this technology and confirm this study’s proof-of-concept design, a viral surrogate of SARS-CoV-2, Human Coronavirus OC43 (HCoV-OC43), was also measured using TOFS. Unlike the highly pathogenic SARS-CoV-2, HCoV-OC43 is the most encountered human coronavirus; it generally causes mild upper-respiratory tract illness and contributes to 15% to 30% of cases of common colds in human adults. Both HCoV-OC43 and SARS-CoV-2 are enveloped beta-coronavirus containing a non-segmented positive-sense, single-stranded ribonucleic acid. The low pathogenicity, high prevalence, and similar density of binding ligand proteins on envelope membranes make HCoV-OC43 an ideal surrogate for label-free direct detection of SARS-CoV-2.

## Materials and Methods

The key element of the present TOFS device is the bitapered optical fiber. Its sizes are illustrated in Figure 1(a). The tapered fiber has a narrow center and waist section with a down-taper and up-taper section on each end. Typical dimensions of TOFS for our experiments are illustrated in Figure 1, with an outer diameter of 125 μm and core diameter of 9 μm optical fiber which only supports single-mode propagation of light. In the tapered region, the outer diameter is 10μm and the core diameter of 0.8 μm. Figure 1(b) shows the fiber with the biomolecular pairs around the taper region, whose concentration is determined through our analysis of the phase change of the optical signal passing through the bitapered fiber.

**Figure 1.**
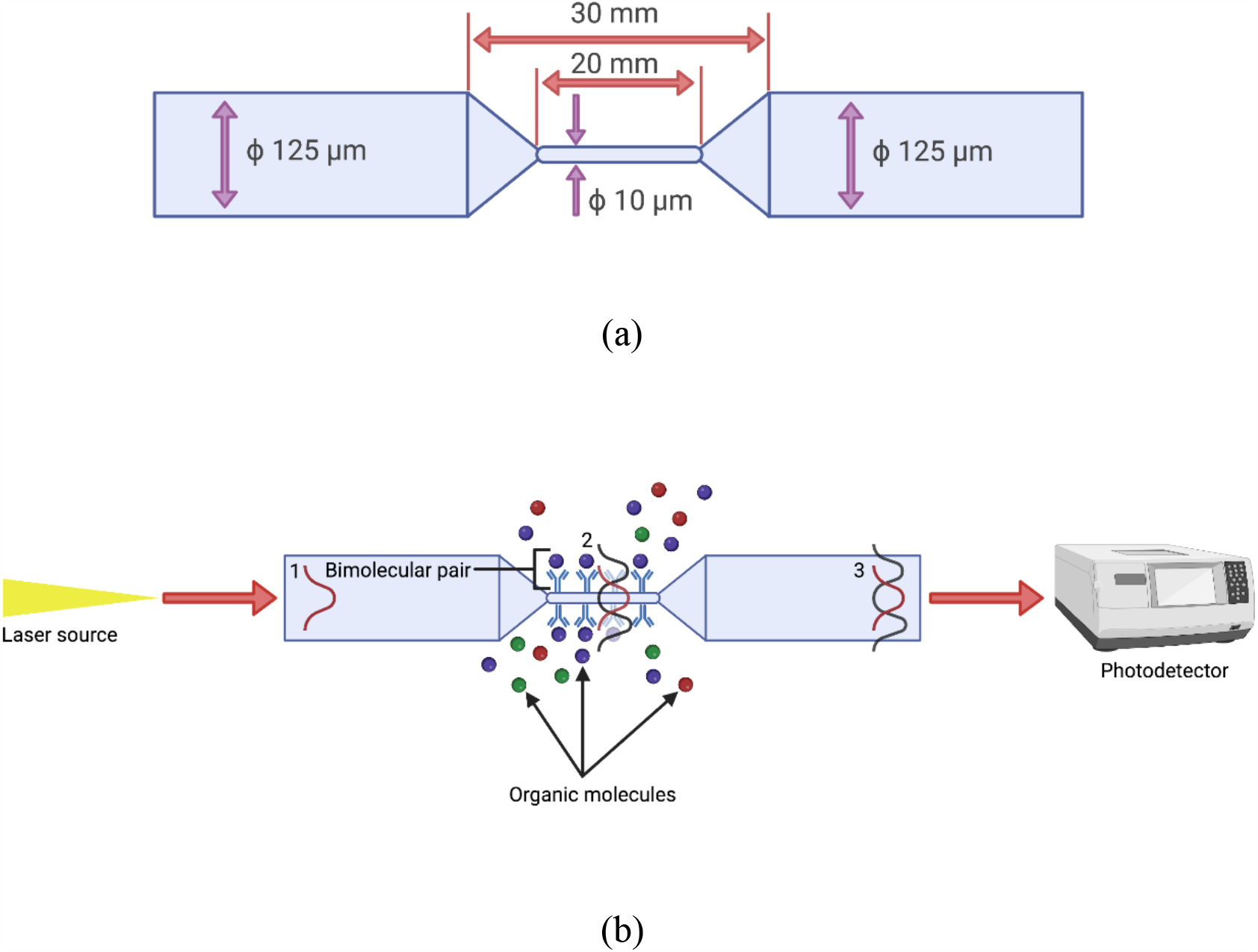
(a) Schematic of typical TOF (not to scale) indicating typical dimensions. (b) The bitapered optical fiber connected to a laser source (input) and detector (output), with the untampered parts 1 and 3, and the taper part where analytes are present.

A photograph of initial laboratory setup used for collecting data is shown in Figure 2(a), with the schematic explained in Figure 2(b). The TOFS system contains three main parts: the tapered optical fiber encased in a reservoir (cell), laser light source, and a photodetector.

**Figure 2.**
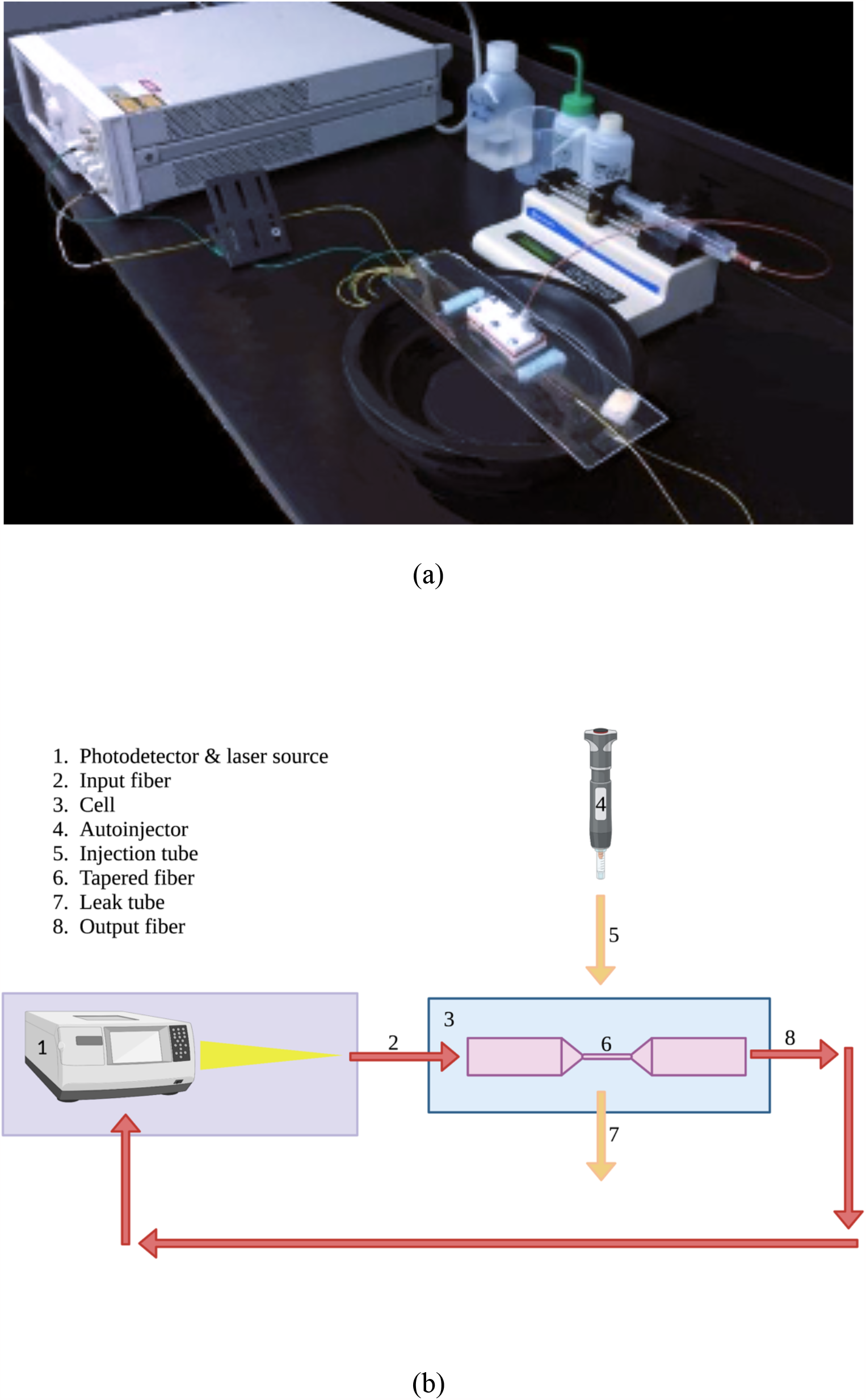
(a) Photograph of the initial system in the lab. After Ref. [Cong et al]. (b) Schematic diagram of the initial laboratory setup.

Light from the source (1) enters the fiber as a single-mode in our standard optical fiber (2) at infrared wavelengths λ ∼1.48 to 1.56 μm. As light passes to tapered region (3), multiple cladding modes are excited simultaneously. These modes have evanescent components in the surrounding medium and are sensitive to the concentration of a bimolecular antigen-antibody pair. These modes revert to a single-mode as they pass through up-taper region (4). The overlap of the modes, either in-phase or out-of-phase with the single-mode, provides an interference spectrum as a function of wavelength. The total transmitted intensity at the output of the fiber (10) is detected by a photodetector (1), where data is analyzed by a virtual interface between the tunable laser and a computer, as shown below. The amplitude and phase of each signal are extracted from the data by Fourier analysis, and the phase change related to maximum amplitude is used.

The introduction of biomolecules or antibodies in the tapered region alters the shape of the oscillations. Analytes may be added to the surrounding waist region of the taper (6) as shown in Figure 2(b), where an autoinjector (4) inserts the material through an injection tube (5) to enter a cell reservoir (3) to alter the environment around the entire taper waist length. Material ultimately drains from the cell via a leak tube (7). The introduction of these biomolecules or antibodies in the tapered region alters refractive index, which is reflected in changes to the signal’s amplitude and phase. The phase change is caused by the change in the effective refractive indexes under different environments, which is altered by the introduction of biomolecules and antibodies. As the concentration of biomolecules and antibodies around the waist region increases, the phase change shifts proportionally.

To achieve high resolution, the signal-to-noise ratio (SNR) must be maximized. This is achieved by optimizing fiber parameters, including the waist diameter and length of the up- and down-taper regions, such that the signal transmits at mainly one dominant frequency.^12^ The signal-to-noise ratio of the system was observed at baseline, prior to analyte introduction, by analyzing the effects of environmental factors on the phase change. Various factors that affect phase change fluctuations include laser detector noise, fiber stress, liquid flow, room temperature, etc. The system requires about 2 hours of warming up to stabilize before data collection. All measurements for one detection were performed in a single day to guarantee consistency.

Optimization of the tapered region dimension has been extensively discussed in Deng *et al*., which was used for this work. The final tapered fiber is placed in a Teflon cell and prepared specifically for label-free detection of the protein IL-8 and the virus HCoV-OC43. Our TOFS is capable of label-free detection due to its ability to detect changes in the refractive index of the tapered fiber surface. This requires the attachment of a molecular detecting element, such as antigen-antibody binding interactions. This allows us to detect target molecules on the femtomolar level. Immunoglobulin G (IgG) antibodies were selected as the recognition element of the tapered fiber. We use silane chemistry to covalently tether IgG molecules to hydroxyl groups on the surface of the tapered fiber. We use the IgG antibody specific for IL-8 protein and the Anti-HCoV OC43 Spike Polyclonal antibody specific for HCoV-OC43 viral antigen in our experiments.^12^ Due to the covalent binding of these antigens to the tapered fiber surface, fibers must be stripped to remove antigens between uses with a 2.4 pH glycine solution.

Recombinant IL-8 was diluted to 8 μg/mL concentration in PBS. PBS rather than saliva was used due to COVID-19 restrictions at the time the experiments were performed. Water, PBS, and saliva have similar refractive indices and our TOFS can use these materials with good signal-to-noise ratios. Therefore, the use of PBS appropriately approximates TOFS detectability of IL-8 dissolved in saliva.

HCoV-OC43 was purchased from ATCC (ATCC® VR-1558) and cultured with human rhabdomyosarcoma cell line RD-151 (ATCC® CCL-136) for acquiring human membrane antigens. Viral titer was determined by indirect immunoperoxidase assay, and the viral particle number was measured with ELISA assays (Using Anti-HCoV OC43 Spike Polyclonal antibody, CABT-CS063, Creative Diagnostics, United States). Anti-HCoV OC43 Spike Polyclonal antibody (CABT-CS063, Creative Diagnostics, United States) was tethered to the tapered fiber as described above. Saliva spiked with HCoV-OC43 (500 μL) was processed with a modified microfluidic filter first developed at the Sinha labs at the University of Southern California.^14^ The microfluidic filter is specially designed with minimal dead volume for filtering viral particles. The RNA-binding member was replaced (as we do not need viral nucleic acid) with a filter membrane of 200-nm pores to allow viral particles to pass while removing cellular debris. The resulting saliva in a syringe was attached to another filter with 80-nm pores. The first 100 μL of saliva passed the filter (without virus as a control input) was added to the reaction chamber for background signal calibration. After removing the control input from the chamber and removing the filter from the syringe, another 100 μL of saliva (from the same sample) with viral particles was added for binding to TOFS (for 5 min). The phase change of light was calculated and processed for HCoV-OC43 detection. This approach yielded reproducible results consistently. The prepared TOFS was then sent back to the University of Dayton.

In practice, the laser took about 40 minutes to stabilize prior to use. Warming the laser up before use allows the laser to stabilize and minimizes system noise.

## Results

### IL-8 detection

The following experiments were conducted with the IL-8 analyte and tapered fiber with mouse anti-human IL-8 antibody (IgG). First, the SNR of the system was measured by calculating the phase change caused by exposure of the tapered fiber to antibodies prior to analyte introduction. Phase shifts from 66 samples related to different conditions were gathered over a span of 6 hours. Details can be found in Cong *et al*.^12^ This is reproduced in Figure 3 with phase changes plotted against the elapsed times. System noise was estimated by the standard deviation of phase change fluctuations between 40 and 70 minutes, a period of laser stabilization. The TOFS system exhibited a noise of approximately 0.0066 radians, significantly less than the phase changes exhibited during periods of IL-8 detection, which ranges from 1.88 to 2.10 radians.

**Figure 3.**
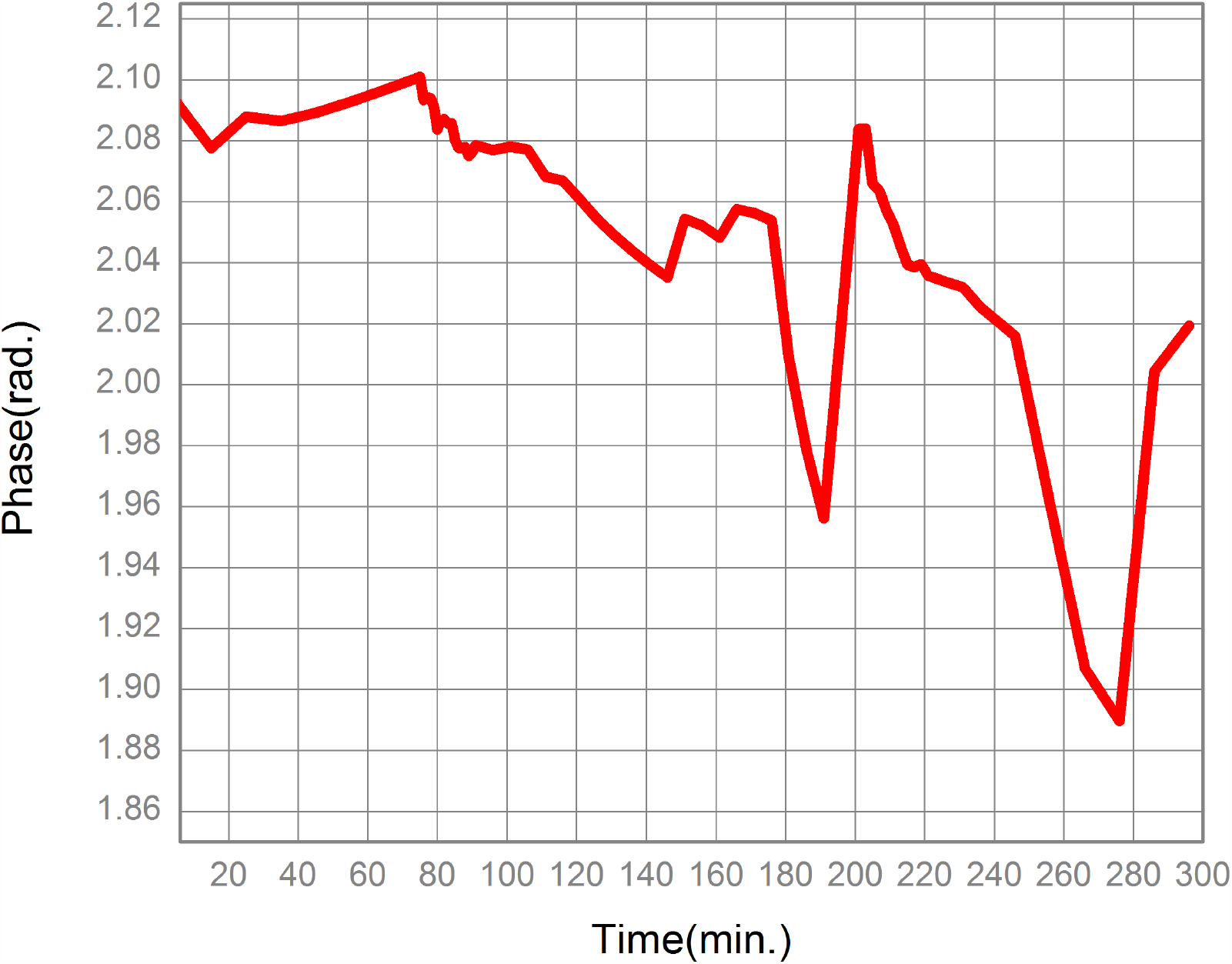
Variation of the signal phase versus real time after fitting from 66 data sets [Cong *et al*.].

We assumed that the amount of IgG required to saturate the tapered fiber is equivalent to the number of molecules that can attach to the tapered fiber surface at any time. Therefore, we can approximate the amount of IgG required to saturate the fiber waist by measuring the ratio of the surface area of the tapered fiber to the surface area of the molecule. We assumed that IgG molecules are 28 nm in diameter and IL-8 molecules are 14 nm [ref]. When IgG bound IL-8 tethered to a TOFB at different concentrations, we observed the expected phase change with a wavelength from 1.48 to 1.56 μm, as reported in Ref.^15^ The two dips, between 170 and 190 min and then between 240 and 290 min, in Figure 3 occur because we repeated this process twice to form antigen-antibody pairing by adding 100 pg/mL at each time. The overall calculated SNR is > 10 for these two drops in the PCs. The TOFS prototype is indeed a very sensitive device for the detection of biomolecules. As little as 10 pg/ml or 7 × 10^5^ IL-8 molecules/μl can be detected by the prototype device, indicating that the sensitivity is better than 10 pg/ml using our sensor.

### HCoV-OC43 detection

To validate our proof-of-concept experiment, a viral surrogate of SARS-CoV-2, Human Coronavirus OC43 (HCoV-OC43), dissolved in saliva was also tested. A typical set of results and its processing, similar to Figure 3, are shown in Figure 4. The intensity spectrum (not shown here) shows that it is significantly different from the corresponding case for IL-8, indicating the possibility that there could be too much antigen-antibody binding that could saturate the SL. However, a dominant frequency can still be identified; this time at 8 instead of 7. A larger dominant frequency (DF) favors more sensitive detection.

**Figure 4.**
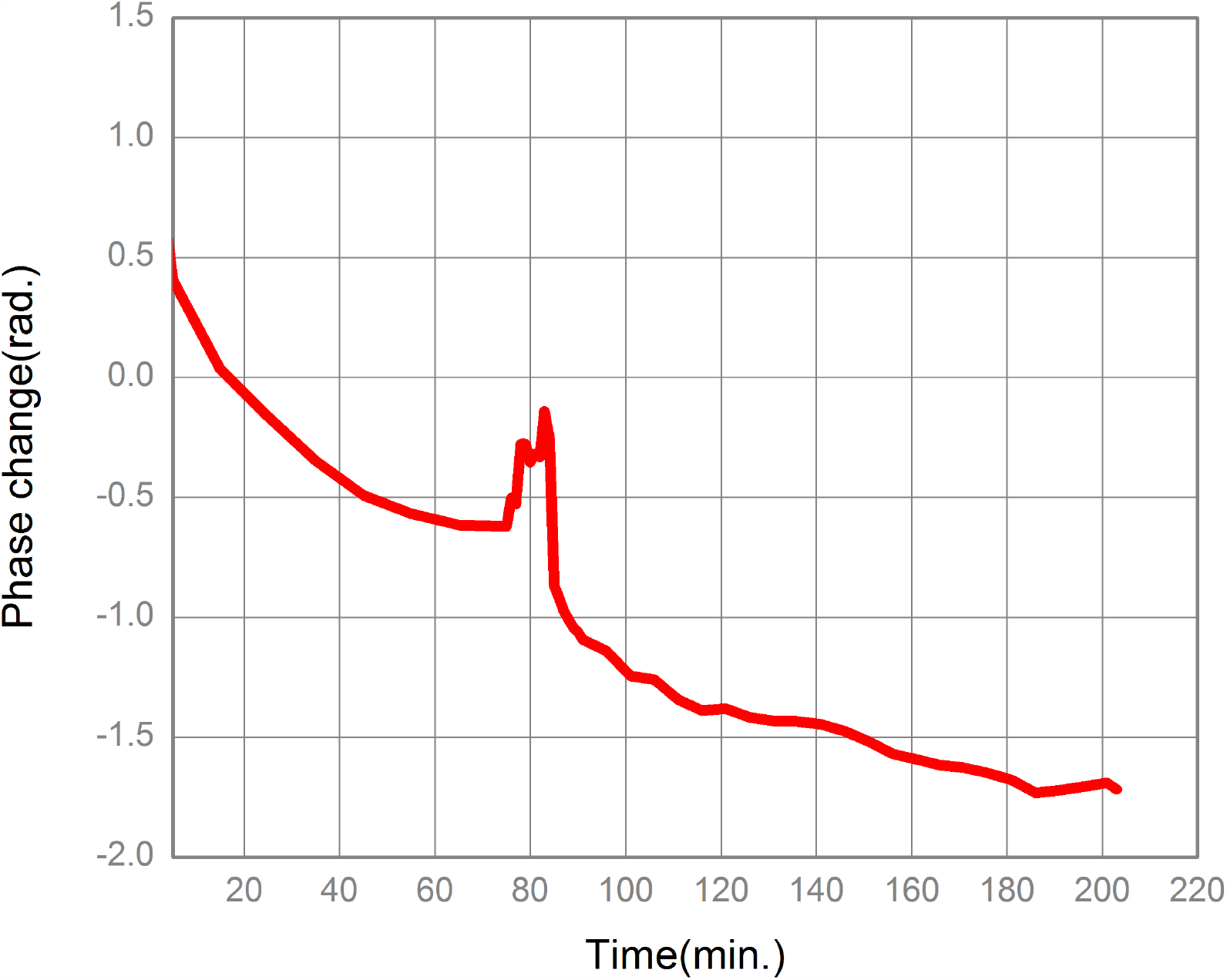
Phase change (PC) signal for HCoV-OC43. 100 µL of antigen was added.

**Figure 4.**
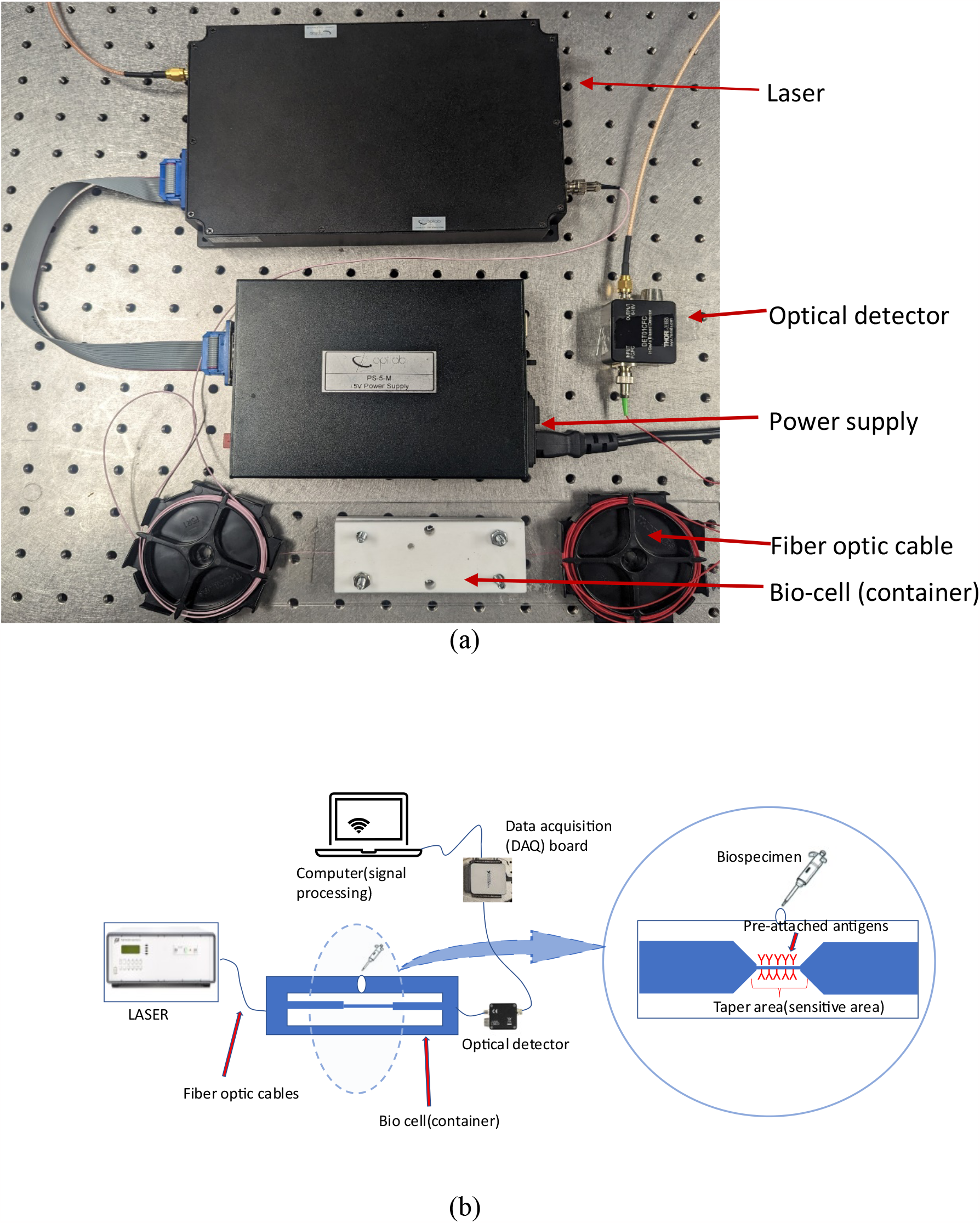
(a) Photograph of latest prototype of TOFS, (b) The schematic diagram of minimized system with new instruments.

### Final prototyping

To make the system smaller than the one shown in Figure 2, a new compact system was implemented as shown in Figure 5(a). It includes three main devices: a frequency (or wavelength) swept laser, a detector, and a data acquisition (DAQ) board. In Figure 5(b), a schematic diagram is illustrated. Its principle of operation is the same as the previous lab setup; the main idea here is to make the system more compact.

**Figure 5.**
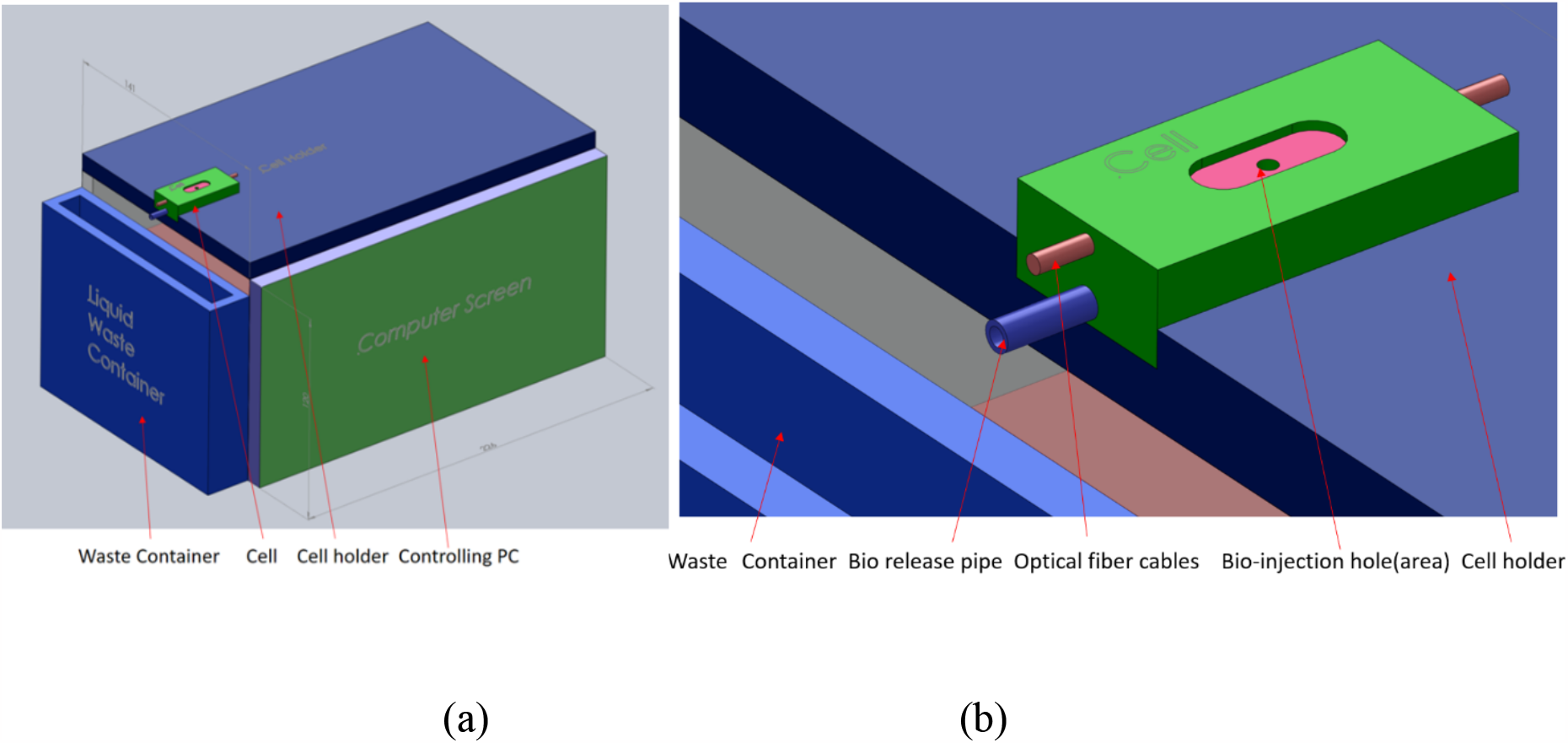
Conceptual device packaging design based on Figure 4. (a) The whole system, (b) zoomed in view of replaceable cell.

In this system the linear relationship of laser wavelength with the externally supplied voltage is used to obtain the required wavelength range between 480-560 nm. The supplied voltage is swept through a certain fiber using a computer-controlled program to obtain above wavelength range. One full sweep of the wavelength scan is considered as one scan of the biosample. We can identify the phase changes when the material around the tapered area is altered (the refractive index around the tapered area). The bio sample is scanned repeatedly in an optimized rate and then a signal processing method is used to analyze the signal output. All these steps are automated, and straightforward for the user. A triangular wave shaped voltage scan was chosen for scanning to avoid abrupt voltage alterations for the laser safety. The sweeping rate is kept at 0.25Hz, or 20.0 nm/s.

The output signal is in the form of phase change when the overall refractive index is changed around the tapered area. This is expected to occur only when reattached antigen material is bonded with any targeted material which would present in a bio sample. If such targeted biomaterial is not present in the bio sample a flat phase graph is expected to receive.

The system was first tested using water. When methanol was added to the cell with water, there was a significant phase change despite that though both methanol and water have similar refractive indices. The results are very consistent and repeatable. This demonstrates the high sensitivity and reliability of the overall system.

Finally, after successful experiments, we designed the packaging all as shown in Figure 5, with all the devices in Figure 4, except for the laptop. It was designed as a compact portable system, in size 226 X 141 X 120 mm, as shown in Figure 5 (a), which was proved to be a practical functional device, supported by many real experiments. This design that can compactly contain all the equipment for easy transportation and to accommodate the cell assembly, as is necessary for point-of-care devices. All functional aspects were considered when the box was designed, viz., heat dissipation of the laser, ease of handling, ease of transportation, and most importantly, protecting the electronics from bioliquids. The disposable cell was also designed and built as shown in Figure 5 (b), with a cost of less than $10 USD.

## Discussion

TOFS is a highly sensitive system capable of detecting proteins, viruses, and other biomolecules. TOFS detected IL-8 with a sensitivity of 10pg/ml or 7.1X10^5^ IL-8 molecules/ml and HCoV-OC43 with a sensitivity of 50 viruses/ml. The noninvasive nature of liquid biopsies such as with TOFs offers clinicians an opportunity for screening not yet achieved in HNSCC, technology that may finally improve HNSCC morbidity and mortality. While a healthy individual has 300 – 500 pg/ml in saliva, HNSCC patients have 1700 – 2500 pg/ml.^14^ Therefore, TOFS sensitivity is more than adequate to detect changes in IL-8 concentration in affected patients, supporting its role in detection and surveillance of HNSCC. The long-term goal for TOFS is its application as a point-of-care device used for detection, monitoring, and surveillance of HNSCC as well as detection of other pathogens in the clinical setting. It has been confirmed that the functional device size can be as small as 141x120x 226 mm. This is very close to a POC or home use unit. Such technology provides the advantage of safe, efficient, portable, and potentially inexpensive results.

Our validation experiment using HCoV-OC43 reinforced the high sensitivity of the system, as TOFS successfully detected the virus with a concentration of 50 viruses/mL. This data clearly supports the use of TOFS in pathogen identification in addition to cancer biomarker detection. Further, this validates the generalizability of the system across various analytes. One such future application involves non-invasive detection of HPV, an oncogenic virus known as a prognostic indicator.^15^ With the increasing prevalence of HPV related HNSCC accounting for about 25% of HNSCC worldwide, salivary HPV may provide a salient screening and prognostic biomarker.^16^ Further, the adaptability of TOFS technology facilitates its application in identifying patients likely to benefit from certain target immunotherapies, such as the detection of PD-1 and PD-L2 biomarkers to predict responsiveness to checkpoint inhibitor therapy.^17,18^ In addition to the utility of TOFS in biomarker detection in the clinic, its application in liquid biopsy has the potential to bolster research in salivary biomarkers of HNSCC.

The reproducibility of TOFs has been confirmed many times for 3 years through multiple fabrications and consistent results. We have also developed a unique discrete fast Fourier transform method for effective real-time data processing with stable and repeatable results, which only selects the dominant harmonic and minimizes the effect of noise. Reproducibility and stability of the system are essential for cancer screening and monitoring due to the high impact of such results. Failure in both could lead to false positives and negatives outside of the systems reported calibrations. False positives can lead to unnecessary imaging and invasive procedures that can be both harmful and costly, in addition to the stress and emotional impact associated with such a work-up. False negatives can result in late disease detection and physician mistrust. The importance of evidence supporting both reproducibility and stability in TOFs cannot be overemphasized.

The long-term goal of the TOFs system is its use as a point-of-care (POC) technology for detection, monitoring, and surveillance in the clinical setting. POC devices must exhibit high sensitivity and specificity, system stability, rapid results, and low cost. As discussed earlier, the present TOFS device has proven its specificity and sensitivity across two analytes and different mediums. Our device is rugged (structurally adiabatic because of the taper), and it has been tested by transporting it multiple times between different locations and with commercial handling. The laser used is stable, with a wavelength sweep of λ ∼1.48 to 1.56 μm. The SNR is 10, which is ample for detecting phase changes. Any fluctuations in laser intensity are accounted for using signal processing as described. Further, it has been determined that changes in the ambient temperature do not significantly affect results, so the system is non-reactive to differences in human temperatures. We have consistently seen and identified the dominant frequency from which the phase is recovered. The LabView program, which is now fully automated, works seamlessly to control the signal generator to supply variable DC voltage values to the laser, collect and store the output data from the cell assembly, and perform signal analysis to calculate the PC values and hence the concentration. Test speed of samples can be achieved in a few seconds.

With the development of a working practical prototype, in the future, we plan to selectively detect target biomolecules in saliva in the presence of other molecules. Our goal is to create a POC device that is amenable to large-scale use in the market with the footprint of a laptop computer and manufacturing on cell assemblies in the price range of a few dollars. We will work on commercialization in the future.

